# Intraspecies genomic divergence of coral algal symbionts shaped by gene duplication

**DOI:** 10.1101/2023.03.28.534646

**Authors:** Sarah Shah, Katherine E. Dougan, Yibi Chen, Debashish Bhattacharya, Cheong Xin Chan

## Abstract

Dinoflagellates of Order Suessiales include the diverse Family Symbiodiniaceae known for their role as essential coral reef symbionts, and the cold-adapted *Polarella glacialis*. These taxa inhabit a broad range of ecological niches and exhibit extensive genomic divergence, although their genomes are in the smaller size ranges (haploid size < 3 Gbp) compared to most other dinoflagellates. Different isolates of a species are known to form symbiosis with distinct hosts and exhibit different regimes of gene expression, but intraspecies whole-genome divergence remains little known. Focusing on three Symbiodiniaceae species (the free-living *Effrenium voratum*, and the symbiotic *Symbiodinium microadriaticum* and *Durusdinium trenchii*) and the free-living outgroup *P. glacialis*, all for which whole-genome data from multiple isolates are available, we assessed intraspecies genomic divergence at sequence and structural levels. Our analysis based on alignment and alignment-free methods revealed greater extent of intraspecies sequence divergence in symbiodiniacean species than in *P. glacialis*. Our results also reveal the implications of gene duplication in generating functional innovation and diversification of Symbiodiniaceae, particularly in *D. trenchii* for which whole-genome duplication was involved. Interestingly, tandem duplication of single-exon genes was found to be more prevalent in genomes of free-living species than in those of symbiotic species. These results in combination demonstrate the remarkable intraspecies genomic divergence in dinoflagellates under the constraint of reduced genome sizes, shaped by genetic duplications and symbiogenesis events during diversification of Symbiodiniaceae.

## Introduction

Dinoflagellates of the Order Suessiales include the Family Symbiodiniaceae, which predominantly consists of symbiotic lineages essential to coral reef organisms. Symbiodiniaceae taxa collectively exhibit a broad spectrum of symbiotic associations (i.e., facultativeness) and variable degrees of host specificity (i.e., host-specialist vs host-generalist), although some are described as solely free-living (Thornhill et al. 2014; LaJeunesse et al. 2018). A comparative analysis of whole-genome sequences from 15 taxa revealed extensive sequence and structural divergence among Symbiodiniaceae taxa, which was more prevalent in isolates of the symbiotic species, *Symbiodinium microadriaticum* (González-Pech et al. 2021). This was supported by a metagenomics survey of single-nucleotide polymorphisms in the genomes of symbiotic *Symbiodinium fitti* from different coral taxa and biogeographical origins, revealing intraspecies sequence divergence correlated to coral host taxa (Reich et al. 2021).

A recent comparative genomic analysis incorporating genomes from three isolates of the free-living species *E. voratum* revealed genome features representative of the Symbiodiniaceae progenitor, due to the absence of symbiogenesis in the *Effrenium* lineage (Shah et al. 2023). These features include longer introns, more extensive RNA editing, less pseudogenisation, and, perhaps most surprisingly, similar genome sizes when compared to symbiotic counterparts. The genome size of *E. voratum* suggests that genome reduction (to haploid genome size < 3Gbp) occurred in symbiodiniacean dinoflagellates before diversification of Order Suessiales (Shah et al. 2023). These results further hint at a role of symbiotic lifestyle in shaping intraspecies genomic divergence and the evolution of these taxa. Intragenomic variation of the ITS2 phylogenetic marker sequences is known among Symbiodiniaceae taxa (Wilkinson et al. 2015; Hume et al. 2019). However, intraspecies whole-genome divergence in these taxa relative to symbiotic versus free-living lifestyle remains little known. Whole-genome data from multiple isolates of a species provide an excellent analysis platform to address this knowledge gap.

Here, we investigate intraspecies genomic divergence in four Suessiales species (of which three are Symbiodiniaceae); these taxa represent two free-living species and two symbiotic species, for which whole-genome data from multiple isolates are available. We focus specifically on sequence and structural conservation, gene family dynamics, and gene duplication, and how these features may reflect adaptation to the distinct lifestyles.

## Results and Discussion

We used four Suessiales species for which multi-isolate genome data are publicly available, to investigate patterns of intraspecies genomic divergence related to facultative lifestyle. The two symbiotic symbiodiniacean species, *S. microadriaticum* (González-Pech et al. 2021; Nand et al. 2021) and *Durusdinium trenchii* (Dougan et al. 2022a), represent taxa that arose from independent origins of symbiogenesis (Figure 1 and Supplementary Table S1). The remaining two are free-living species, the symbiodiniacean *E. voratum* (Shah et al. 2023) and *Polarella glacialis* that is sister to the Symbiodiniaceae in the Order Suessiales (Stephens et al. 2020). The available genome data were generated from isolates collected over vast geographic areas: the thermotolerant symbiont *D. trenchii* from the Caribbean Sea and Pacific Ocean, the free-living *E. voratum* from the Mediterranean Sea and both sides of the Pacific Ocean, the symbiotic *S. microadriaticum* from the Red Sea, Pacific Ocean, and the Caribbean Sea, and the psychrophilic *P. glacialis* from the Antarctic and Arctic oceans (Figure 1). Collectively, these data provide a robust analytic framework for interrogating intraspecies genomic divergence.

**Figure 1.**
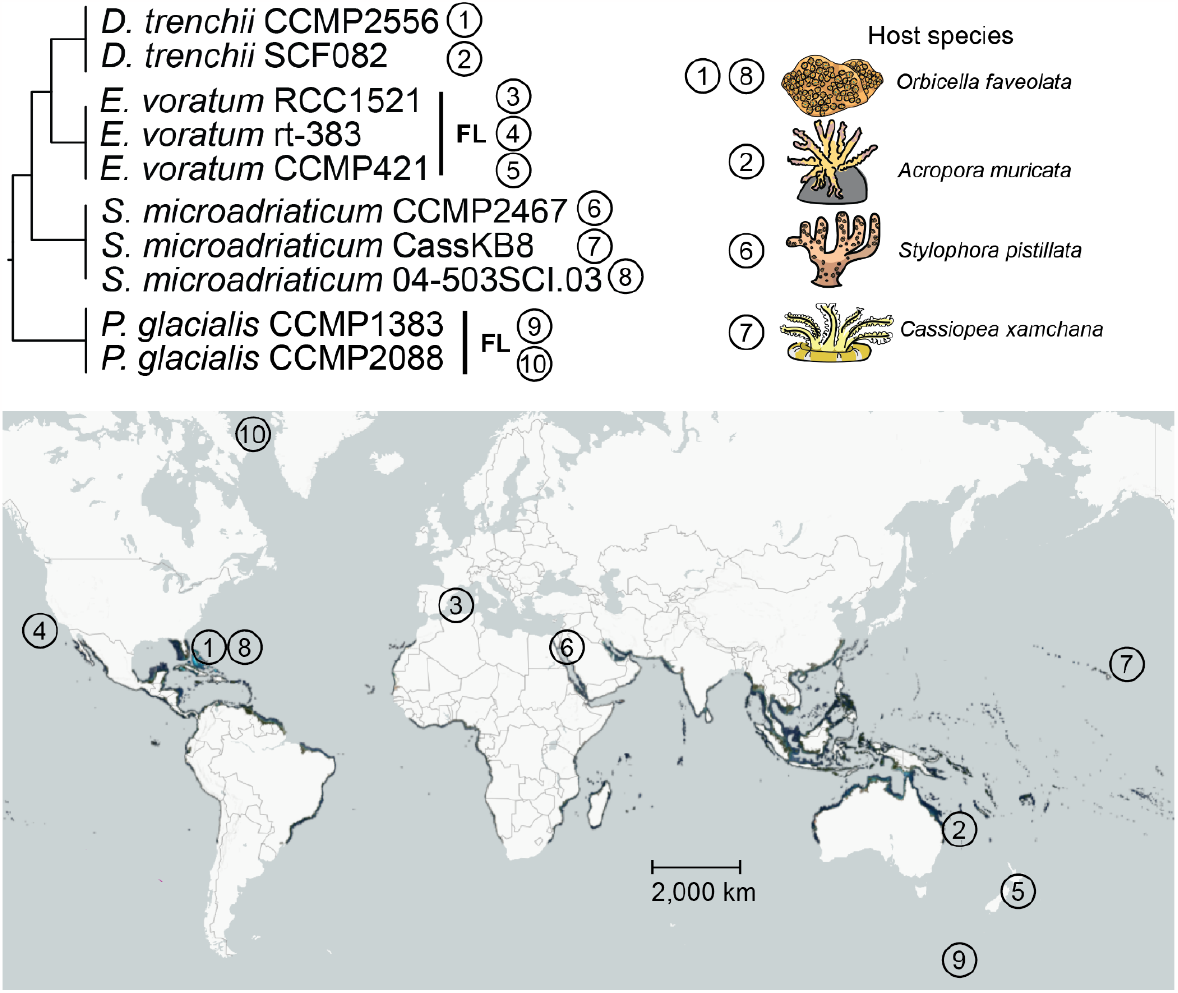
Suessiales species, following LSU rDNA phylogeny (LaJeunesse et al. 2018), for which genome data of multiple isolates are available. Coral reef (in dark blue and cyan) world map by Allen Coral Atlas (2022). Those not marked FL (free-living) are symbiotic and their host species are represented on the top right.

### Genomes of facultative symbionts exhibit higher sequence divergence

We investigated divergence of genome sequence following the approach of González-Pech et al. (2021). For each pairwise comparison of genome sequences, we calculated the percentage of aligned bases, *Q*, and overall sequence identity of aligned regions, *ID*. Genome sequences from isolates of the same species are highly similar (*Q* > 70.2%, *ID* > 98.6% with minimum alignment length 100 bp; Figure 2A, see Supplementary Figure S1 for detail), compared to those between species (*Q* < 10.0%, *ID* < 98.6%). High intraspecies sequence similarity was observed despite the diverse geographic origins for isolates from each species (Figure 1). Genome sequences of the free-living *P. glacialis* were the most similar (*Q* = 95.5%, *ID* = 98.7%; CCMP1383 against CCMP2088), followed by the symbiotic *D. trenchii* (*Q =* 93.3%, *ID* = 99.8; CCMP2556 against SCF082), the free-living *E. voratum* (*Q* = 92.0%, *ID* = 99.4%; RCC1521 against rt-383), and the symbiotic *S. microadriaticum* (*Q* = 78.5%, *ID =* 99.7%; CCMP2467 against CassKB8). Among the three *E. voratum* isolates, CCMP421 showed smaller percentage of aligned genome bases against rt-383 (*Q* = 70.2%) and against RCC1521 (*Q* = 79.2%), compared to *Q* = 92.0% observed between RCC1521 and rt383; this is likely due to the more-fragmented CCMP421 genome assembly, also reflected in the low percentage of mapped sequence reads (Supplementary Table S2). Between the two symbiotic species, the greater divergence observed in *S. microadriaticum* might represent its much earlier emergence and diversification (LaJeunesse et al. 2018). Alternatively, the lower divergence in *D. trenchii* may be due to the recent whole-genome duplication (WGD) in this lineage (Dougan et al. 2022a). Genome data of multiple isolates from a broader taxon representation of Symbiodiniaceae lineages will help clarify the possible link between intraspecies divergence and facultative lifestyle of these symbionts.

**Figure 2.**
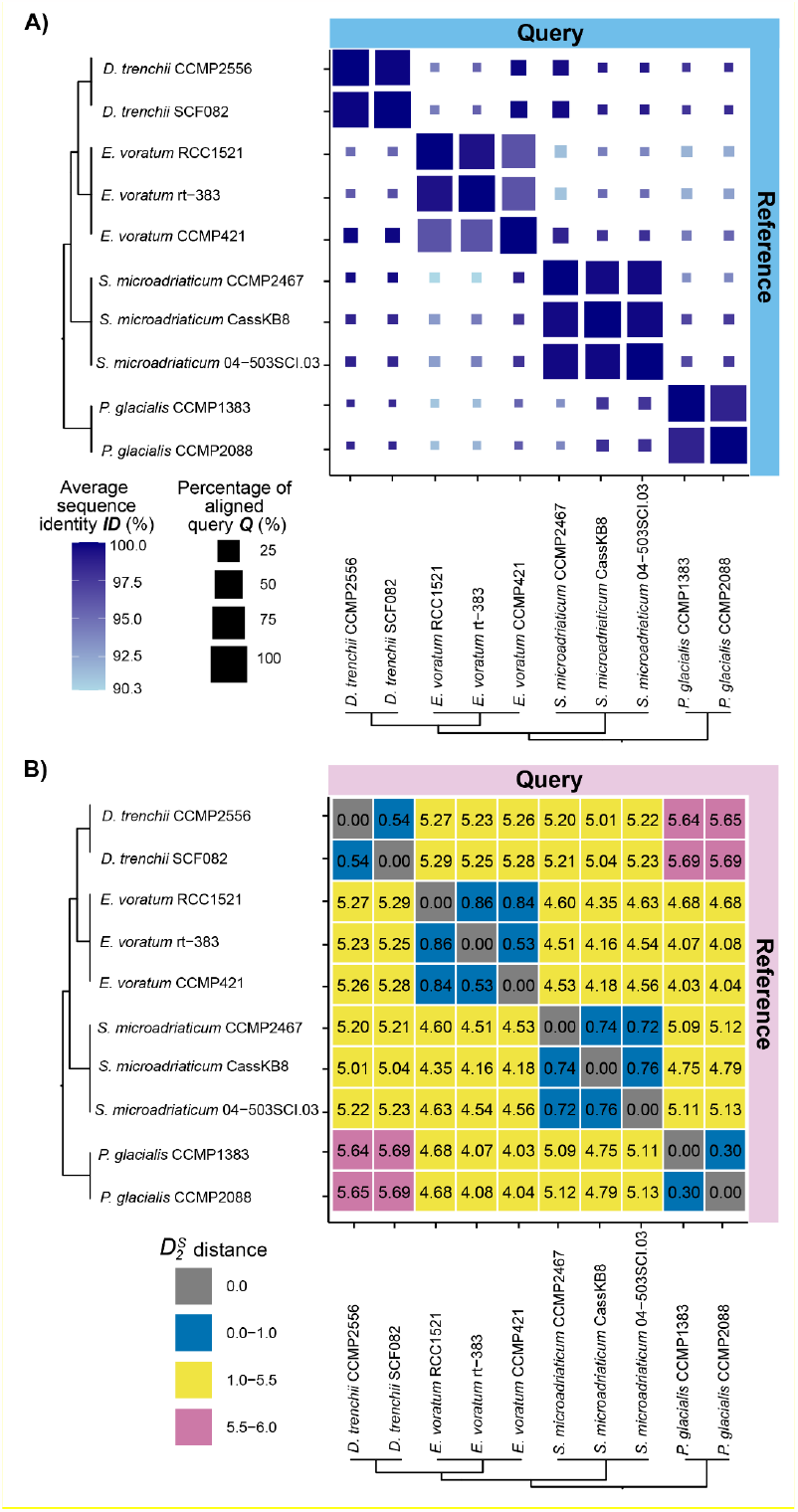
Intra/interspecies genome sequence identity among the four Suessiales species. (A) Alignment-based identity (minimum alignment length = 100 bp) with query genome sequences (y-axis) aligned to the references (x-axis). The colour of the squares corresponds to percent sequence identity *ID* (darker blue = higher identity) and the sizes represent the percentage of the query genome sequence *Q* aligned to the reference. (B) Alignment-free 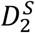 distances (*d*) showing delineation between species (*d* < 1 in blue), Family (*d* between 1.0 and 5.5 in yellow), and the longest evolutionary distance across the Order (*d* > 5.5 in pink).

To extend genome comparisons beyond alignable sequence regions, we further assessed sequence divergence using an alignment-free *k*-mer-based approach. This approach was found to be robust against the contiguity of genome assemblies (Dougan et al. 2022c), and has been applied successfully to discover distinct phylogenetic signals in different genomic regions of Symbiodiniaceae (Lo et al. 2022; Shah et al. 2023). We followed Lo et al. (2022) to derive pairwise 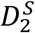 distances, *d*, based on shared *k*-mer profiles at *k =* 23 observed in whole-genome sequences (see Methods). As shown in Figure 2B, the lowest sequence divergence was seen in *P. glacialis* (*d* = 0.30), followed by *E. voratum* (*d* = 0.53 between RCC1521 and rt-383; *d* = 0.9 when implicating the more-fragmented CCMP421 assembly), *D. trenchii* (0.54), and the three *S. microadriaticum* isolates (0.72-0.76). This pattern of divergence is consistent with our observations based on *Q* and *ID* in Figure 2A.

We further assessed the conserved core 23-mers in each species (i.e., *k*-mers common in genomes of all isolates within a species). For each species, we assessed the extent of genome content shared among the isolates based on *x*, the percentage of core 23-mers relative to all distinct 23-mers; in the perfect scenario where genomes of all isolates are identical, *x* = 100%. Using this approach, *E. voratum* and *S. microadriaticum* show similar extent of shared genome content among their corresponding isolates (*x* ranges between 19.5% and 25.2%; Supplementary Table S3). Approximately two-fold greater *x* was observed for *P. glacialis* (52.3-54.9%) and *D. trenchii* (55.6-55.7%); this observation likely reflects the impact of a diploid genome assembly in the former (Stephens et al. 2020) and WGD in the latter (Dougan et al. 2022a). Duplicated genomic regions arising from WGD are resolved over long evolutionary time scales of hundreds of millions of years (Carretero-Paulet and Van de Peer 2020). Given the recent (∼1 MYA) WGD in *D. trenchii*, this species likely has not had sufficient time to resolve genetic redundancy. Regardless, our results here lend support to the general utility of *k*-mer-derived distances in clarifying genome-sequence divergence beyond gene boundaries, which may serve as evidence to guide or complement taxonomic classification of Symbiodiniaceae, and potentially of other dinoflagellates (Dougan et al. 2022c).

### Intraspecies structural divergence in the genomes of Symbiodiniaceae

To assess intraspecies structural genomic divergence, we identified collinear gene blocks in all possible pairwise genome comparisons for each species (see Methods); the greater recovery of these blocks and their implicated genes indicates a greater conserved synteny among the isolates in a species. As expected, due to recent WGD, the two symbiotic *D. trenchii* isolates CCMP2556 and SCF082 displayed the greatest conserved synteny (1,613 blocks implicating ∼22% of total genes spanning 181-199 Mbp; Supplementary Table S4). On the other hand, genomes of the symbiotic *S. microadriaticum* (100-196 blocks, 2.7-3.6% of genes, 8.1-16 Mbp) showed less conserved synteny than the free-living *E. voratum* RCC1521 and rt383 (344 blocks, 6.6-8.1% of genes, 51-60 Mbp; Supplementary Table S4); at first glance this result appears to support observations in an earlier study (González-Pech et al. 2021) that the extent of structural rearrangements is greater in genomes of facultative symbionts than those of free-living taxa. However, the greater contiguity of the *E. voratum* assemblies (scaffold N50 length = 720 Kbp for RCC1521, 252 Kbp for rt-383) than that of *S. microadriaticum* assemblies (e.g., scaffold N50 length = 43 Kbp for CassKB8 and 50 Kbp for 04-503SCI.03) represents a systematic bias that would affect recovery of collinear gene blocks. *S. microadriaticum* CCMP2467 (N50 length 9.96 Mbp) (Supplementary Table S1), the sole representation of a chromosome-level assembly, lacks comparative power in this instance. As a case in point, the inclusion of the fragmented assembly of *E. voratum* CCMP421 (N50 length 304 Kbp; 38,022 scaffolds) lowers the extent of conserved synteny identified in *E. voratum* (195-331 blocks, 4.4-7.9% of genes, 30-65 Mbp; Supplementary Table S4), and we identified no collinear gene blocks between the outgroup *P. glacialis* isolates due in part to sparsity of genes on the assembled genome scaffolds (Stephens et al. 2020). These results in combination suggest that while structural rearrangements contribute to structural divergence of Symbiodiniaceae genomes as postulated in those of facultative symbionts (González-Pech et al. 2019) even at intraspecies level, such an analysis based on collinear gene blocks is sensitive to contiguity of assembled genome sequences. An in-depth assessment of structural divergence would require genome assemblies of comparably high quality.

### Genetic duplication enables functional innovation

We assessed the evolution of protein families for evidence of functional innovation and divergence within species, and its relation to lifestyle. For each species, we inferred homologous protein sets with OrthoFinder using sequences predicted from all corresponding isolates (see Methods); the homologous sets that are specific to an isolate may reflect instances of contrasting divergence in and/or specialisation of protein functions (e.g., putative remote homologs), occurring at distinct evolutionary rates. First, we assessed number of isolate-specific sets for each species based on OrthoFinder results ran at default parameters (i.e., inflation parameter *I* = 1.5). The highest percentage of isolate-specific sets was observed in *D. trenchii* (17.2% of total sets), followed by *P. glacialis* (16.0%); these numbers are nearly four-fold greater than that observed in *S. microadriaticum* (4.0%) and *E. voratum* (4.1%; Figure 3). To investigate the robustness of this result, we increased the inflation parameter (*I*) for clustering within OrthoFinder that controls the granularity (i.e., higher inflation parameter produces smaller clusters). As expected in all cases, the increase of *I* resulted in an increase of isolate-specific protein sets; at *I* = 10, the percentage of these sets is 37.8% (*D. trenchii*), 32.4% (*P. glacialis*), 15.6% (*S. microadriaticum*), and 10.8% (*E. voratum*). Despite the high synteny and sequence conservation in *D. trenchii*, the substantial number of protein families retained in duplicate after WGD show evidence of isolate-specific divergence and/or specialization in *D. trenchii* where facultative lifestyle has been hypothesized to be the main driver of post-WGD adaptation (Dougan et al. 2022a). On the other hand, the comparable extent of isolate-specific protein sets in *P. glacialis* may represent heterozygosity inherent to a diploid representation of the genome assembly (Stephens et al. 2020), distinct from the haploid genome assemblies among the Symbiodiniaceae taxa. None of the *E. voratum* and *S. microadriaticum* isolates showed evidence of WGD (Supplementary Table S5), and thus the similar level of isolate-specific divergence in these species supports the notion of massive genome reduction in the Suessiales ancestor, with WGD a mechanism for escaping this process to generate functional innovation, as observed in *D. trenchii* (Dougan et al. 2022a).

**Figure 3.**
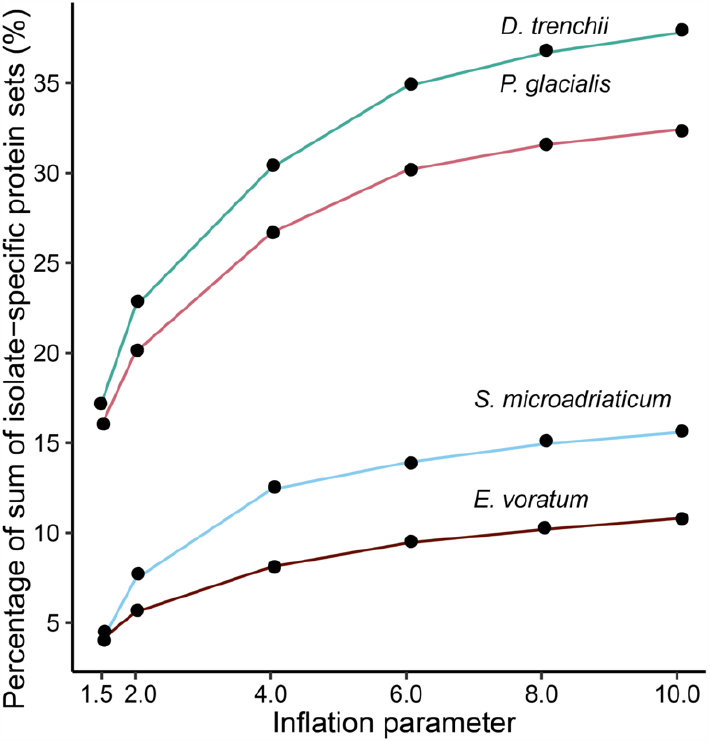
The percentage of isolate-specific protein sets in each Suessiales species. Protein sequences were clustered at inflation parameter *I* between 1.5 and 10 using OrthoFinder.

### Genomes of free-living species exhibit greater extent of tandemly duplicated single-exon genes

Tandemly duplicated (TD) genes, i.e., duplicated genes found next to each other on the genome, are part of unidirectional gene clusters commonly found in dinoflagellates, thought to facilitate their expression (Nand et al. 2021; Chen et al. 2022). In an earlier study (Stephens et al. 2020), ∼40% of the gene repertoire in *P. glacialis* genomes were located in unidirectional gene clusters, many of which encoded functions associated with cold and low-light adaptation. Here we defined a TD block as a block comprising two or more consecutive genes with high sequence identity on a genome scaffold (see Methods). In our independent survey of TD genes in all 19 available Suessiales genomes, we found the largest number and proportion of TD genes in the free-living lineages of *P. glacialis* (7.8% in CCMP1383, 9.2% in CCMP2088) and *S. natans* (7.1%), followed by the symbiotic *S. tridacnidorum* CCMP2592 (6.5%) and *C. goreaui* SCF055 (6.0%), with smaller proportions observed in the free-living *E. voratum* (3.9% in rt-383, 4.4% in RCC1521), and the smallest in *S. microadriaticum* (1.0-2.2%) (Table 1). Some of the largest TD blocks consisted of 13-16 genes, found in genomes of free-living lineages (*S. natans*, and the *P. glacialis* CCMP1383 and CCMP2088). Among the free-living *E. voratum* isolates, the TD block sizes were slightly smaller, implicating genes encoding ribulose bisphosphate carboxylase (the largest block of 9 genes in RCC1521), HECT and RLD domain-containing E3 ubiquitin protein ligase 4 (rt-383, 7 genes), calmodulin (rt-383, 7 genes), and solute carrier family 4 (rt-383, 7 genes) (Supplementary Table S6); these implicated functions are essential for photosynthesis, ion binding, and transmembrane transport. However, we cannot dismiss the possibility of genome-assembly contiguity in affecting recovery of TD blocks. For instance, the recovery of TD genes in the chromosome-level assembly of *S. microadriaticum* CCMP2467 is 2.2% versus ∼1.0% in the other two assemblies, and the recovery of 1.5% in *E. voratum* CCMP421 contrasts to 3.9-4.4% in the other two *E. voratum* genomes. Despite this, a greater extent of TD genes in free-living lineages (*P. glacialis*: 55.2-59.4%; *E. voratum* RCC1521: 23.1% and rt-383: 22.5%; *S. natans*: 21.8%) were single-exon genes, in contrast to the symbiotic *D. trenchii* and *S. microadriaticum* (4.2-9.2%) (Table 1). Our results lend support to the notion that tandem duplication may facilitate transcription of genes encoding essential functions implicating single-exon genes, and is potentially more prominent in genomes of free-living taxa than those of symbiotic lineages (Stephens et al. 2020).

**Table 1.**
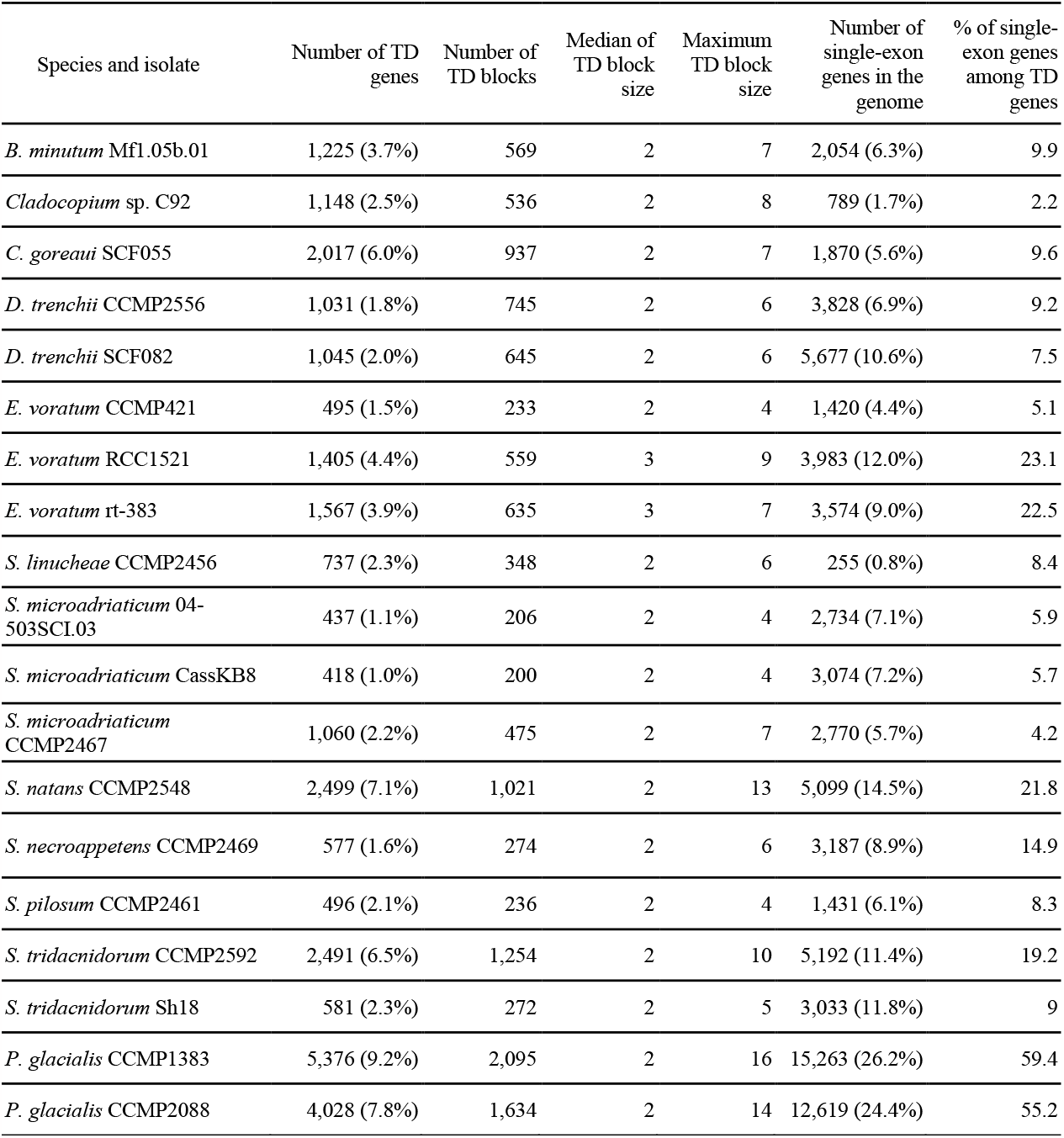
Tandemly duplicated (TD) genes within 19 Suessiales isolates. TD genes were defined as ≥ 2 consecutive genes on the same scaffold making up a “block”, with its size represented by the total number of consecutive TD genes.

Introner elements (IE) are non-autonomous mobile elements characterised by inverted repeat motifs within introns that are hypothesised to propagate introns into genes (Worden et al. 2009; van der Burgt et al. 2012; Huff et al. 2016), which have been found to be more prevalent in genomes of free-living dinoflagellate species (Farhat et al. 2021; Dougan et al. 2022b; Shah et al. 2023). We examined the presence of these elements in the assembled genomes and TD genes for the multi-isolate Suessiales species (Supplementary Table 1). We found the proportion of IE-containing genes overall to be less in Symbiodiniaceae (3.2-6.3%) than *P. glacialis* (10.7-11.5%), a trend also observed in the genome of bloom-forming dinoflagellate species, *Prorocentrum cordatum* (10.4%) (Dougan et al. 2022b). Nonetheless, IEs were only found in a small proportion of TD genes (2.5-5.7%) per Suessiales isolate, suggesting they are neither connected to lifestyle nor play a major role in propagating TD genes in Suessiales (Supplementary Table S1).

### Most tandemly duplicated genes undergo purifying selection

To assess selection acting on TD genes, we focused on the two best-quality genome assemblies (based on number of scaffolds and N50 length) from each species (i.e., total of eight isolates), excluding the fragmented assemblies of *E. voratum* CCMP421 and *S. microadriaticum* CassKB8. We calculated the ratio *ω* as the nonsynonymous substitution rate (*K*_*a*_) to synonymous substitution rate (*K*_*s*_) between all possible gene pairs within each TD block (Supplementary Table S6; see Methods); in general, *ω* > 1.0 indicates positive selection, *ω* = 1.0 indicates neutral selection, whereas *ω* < 1.0 indicates purifying selection (Yang and Bielawski 2000) among TD genes within a block. Based on this analysis, compared to genomes of symbiotic species, those of free-living species yielded larger proportions of TD blocks with mean *ω* < 1.0, indicating purifying selection, i.e., 71.7% in *P. glacialis* and 67.7% in *E. voratum*, compared to 64.2% in *D. trenchii* and 49.1% in *S. microadriaticum* (Figure 4A; Supplementary Table S7). In all cases, the mean *K*_*s*_ value per TD block is less than 0.5 (Figure 4B). The observed mean *ω* values are similar between two isolates of a species, e.g., mean variance of *ω* = 0.26 for both *P. glacialis* isolates (Supplementary Figure S2), suggesting a common pattern of selective pressures acting on TD genes for the species. An exception is the symbiotic *S. microadriaticum* (mean variance of *ω* = 0.16 for 04-503SCI.03 and 0.95 for CCMP2467; Supplementary Figure S2), but more genome data from other multi-isolate symbiotic species will enable the systematic investigation of the possible links between selection acting on TD genes and lifestyles.

**Figure 4.**
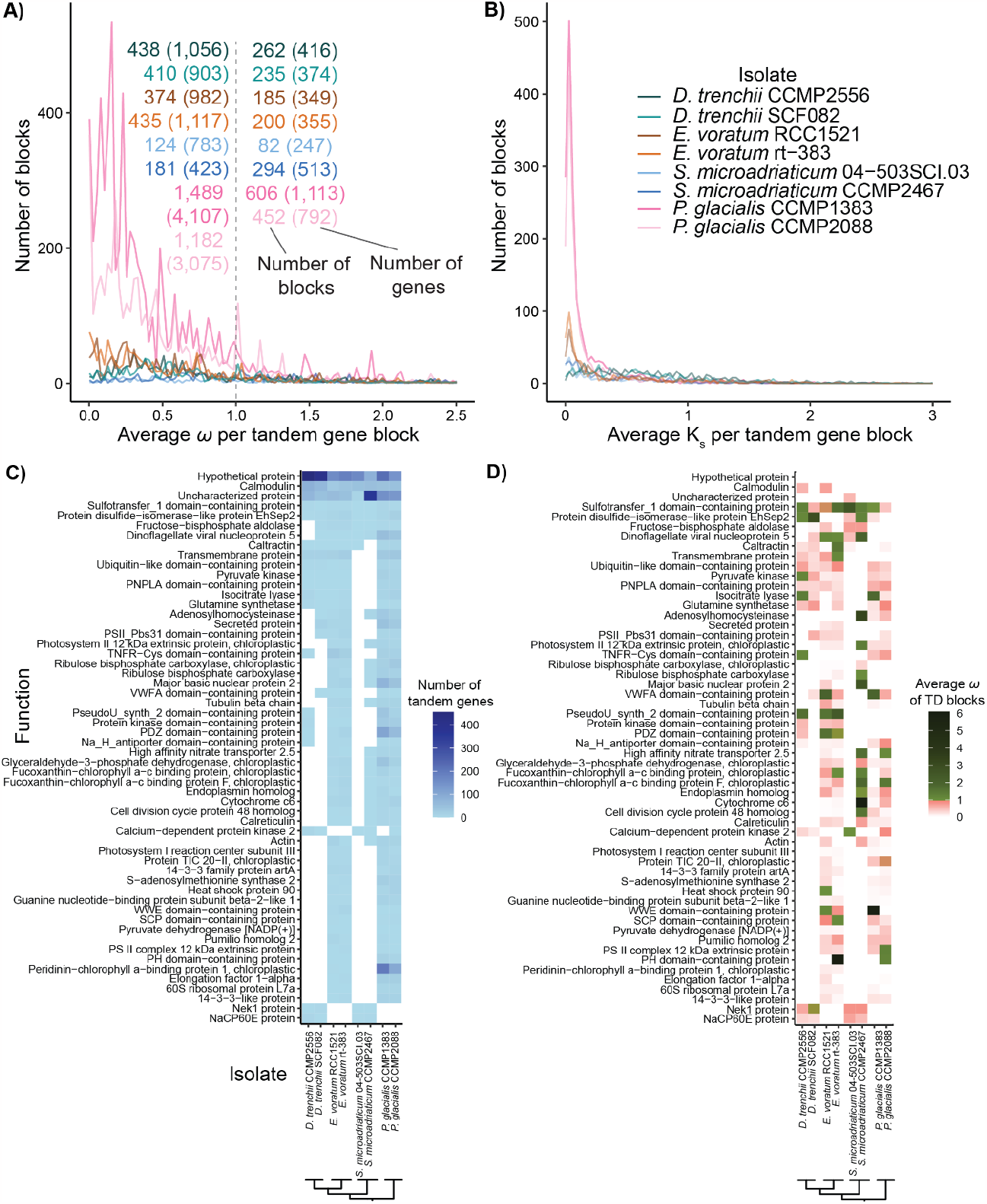
TD genes and their functions in eight Suessiales isolates. The number of TD blocks showing distribution respectively for (A) mean *ω* and (B) mean *K*_*s*_ of each TD block and its associated TD genes with *ω* < 1 or > 1. Functions encoded by TD blocks that were recovered in genomes of both isolates in one or more species, showing the (C) sum of TD genes, (D) mean *ω*.

To assess functions encoded by TD genes, we focused on TD gene blocks that were recovered in genomes of both isolates in one or more species. Functional annotation of these gene blocks is shown in Figure 4C, and the mean *ω* value for the corresponding block is shown in Figure 4D. Genes encoding calmodulin, sulfotransfer domain-containing proteins, and disulfide-isomerase proteins were recovered in TD blocks in all eight isolates. Fructose-bisphosphate aldolase, dinoflagellate viral nucleoproteins, and caltractin were recovered in at least 7 of the 8 isolates. Genes in TD blocks recovered only in free-living *P. glacialis* and *E. voratum* encode functions related to photosynthesis (i.e., photosystem I reaction centre subunit III, chloroplast TIC 20-II protein, PS II complex 12 kDA extrinsic protein, and peridinin-chlorophyll *a*-binding protein). In comparison, those in TD blocks found only in the two symbiotic species encode for Nek1 protein that is involved in maintaining centrosomes, and NaCP60E, a sodium channel protein. Most of these functions were encoded by no more than 50 TD genes per isolate (Figure 4C) in which the mean *ω* per gene block was < 1 (Figure 4D). These results do not speak directly to the specificity of gene functions to tandem duplication in the genomes we analysed, given that some gene copies may also occur elsewhere in the genomes. However, our results suggest a tendency for TD genes within a block to undergo purifying selection, regardless of lifestyle.

### Concluding remarks

Our results, based on multi-isolate whole-genome data from representative species, demonstrate how facultative lifestyle or the lack thereof has shaped the genome evolution of Symbiodiniaceae dinoflagellates. Generation of genetic and functional diversity at the intraspecies level implicates genetic duplication, including tandem duplication of genes. All these evolutionary regimes are under the constraint of genome reduction that is hypothesised to pre-date the diversification of Order Suessiales (Shah et al. 2023). Although our results hint at the potential linkages of facultative lifestyles to some of the varying features observed between free-living versus symbiotic species, whole-genome data from a broader taxonomic representation (and from multiple isolates) will enable a more-systematic investigation to establish these linkages.

## Methods

### Data

For this study, we used publicly available genome assemblies and gene models of *D. trenchii* CCMP2556 and SCF082 (Dougan et al. 2022a), *E. voratum* isolates RCC1521, rt-383, and CCMP421 (Shah et al. 2023), *S. microadriaticum* CCMP2467 (Nand et al. 2021), 04-503SCI.03 and CassKB8 (González-Pech et al. 2021), and *P. glacialis* CCMP1383 and CCMP2088 (Stephens et al. 2020) (Supplementary Table S1). To contrast the contiguity of these genome assemblies, we obtained chromosome numbers from cytological observations (Blank and Trench 1985; Jeong et al. 2014; Wham et al. 2017). For tandem gene duplication analysis, we used genomic datasets from 9 more Symbiodiniaceae isolates (Supplementary Table S1) generated in Chen et al. (2020; 2022), González-Pech et al. (2021), and Shoguchi et al. (2013; 2018). To determine the intraspecific identity of the three *E. voratum* genome datasets, we mapped the short-read gDNA of each isolate obtained from (Shah et al. 2023) to each other using Bowtie2 v2.4.4 (Langmead and Salzberg 2012) with the *--very-fast* algorithm.

### Assessment of genome-sequence similarity based on alignment

To assess genome-sequence similarity of the four target species based on sequence alignment, we used nucmer (*--mum*) implemented in MUMmer 4.0.0beta2 (Marçais et al. 2018) at minimum alignment lengths of 100 bp, 1 Kb, and 10 Kb to align assembled genome sequences for every possible pair of isolates in each species. For each pairwise comparison, we calculated the percentage of aligned bases, *Q*, and overall sequence identity of aligned regions, *ID*. Maximum values of for both *Q* and *ID* at 100% indicate that two genome sequences are identical. We then used mummerplot (*-f --layout*) and dnadiff to generate figures and reports for these alignments.

### Assessment of genome-sequence similarity using an alignment-free approach

Adopting the same approach described in Lo et al. (2022), we calculated 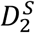 statistic based on shared *k*-mers for each pair of genomes, from which a distance (*d*) was derived. Briefly, Jellyfish v2.3.0 (Marçais and Kingsford 2011) was used to derive *k-*mers (at *k* = 23) from each genome assembly, from which distances were calculated using *d2ssect* (https://github.com/bakeronit/d2ssect) from all possible pairs of genomes. Following the earlier studies (Lo et al. 2022; Shah et al. 2023), core 23*-*mers among isolates of each species were identified from the extracted 23-mers, using the bash command *comm* (−12). BEDtools (Quinlan and Hall 2010) *intersect* was used to find regions of overlap between the core *k-*mers and different genomic features.

### Gene family evolution and introner element search

To infer homologous protein sets among isolates for a species, all protein sequences predicted from all isolates were used as input for OrthoFinder v2.5.4 (Emms and Kelly 2019). The analysis was conducted at different inflation parameters (*I* = 1.5, 2.0, 4.0, 6.0, 8.0, or 10.0). From the generated homologous protein sets, the proportion of isolate-specific sets was identified. To identify introner elements, we used the introner element sequences identified in Shah et al. (2023) from eight Suessiales isolates as a reference for Pattern Locator (Mrázek and Xie 2006) to search for inverted and direct repeat motifs within introns.

### Identification of collinear gene blocks and types of gene duplication

To identify collinear gene blocks shared by isolates of a species, we first identified homologous protein sequences using BLASTp (e-value < 10^−5^, query or subject cover > 50%, filtered for top five hits for each query). This output was used as input for MCScanX (Wang et al. 2012) (*-b 2*) to search for collinear gene blocks between all possible pairs of isolates. For *D. trenchii*, we filtered out duplicated genes (Dougan et al. 2022a) from the MCScanX output by selecting gene pairs that were more similar to each other (i.e., low nonsynonymous (*K*_*a*_) + synonymous (*K*_*s*_) substitution score), then chose gene blocks that still contained ≥ 5 genes. Gene Ontology (GO) terms were assigned to all gene sets via UniProt (version 2022_01) to GO (version December 2022) ID mapping on the UniProt website (uniprot.org/id-mapping). The *duplicate_gene_classifer* implemented in MCScanX was used to assess five distinct type of gene duplications: 1) singleton = not duplicated, 2) dispersed = duplicated with > 10 genes in between, 3) proximal = duplicated with < 10 genes in between, 4) WGD = whole or segmental genome duplication inferred by anchor genes in collinear gene blocks comprising at least 5 genes, 5) tandem = duplicated one after the other, i.e., two or more consecutive genes on the same scaffold.

### Analysis of tandemly duplicated genes

Tandemly duplicated (TD) genes were identified based on the results of MCScanX above. For this analysis, we focused on two best-quality genome assemblies from each species, i.e., for a total of eight genomes. For each TD block, we calculated the nonsynonymous substitution rate (*K*_*a*_) and synonymous rate (*K*_*s*_) between all possible pairs of genes within the block, using the *add_ka_and_ks_to_collinearity*.*pl* script implemented in MCScanX (Wang et al. 2012). The ratio *ω* was defined as *K*_*a*_/*K*_*s*_. When assessing mean *ω* for each TD block, instances of infinity values, e.g., due to *K*_*s*_ = 0, were ignored.

## Supporting information

Supplementary Tables S1 through S7

## Author contributions

Conceptualization, SS, KED, DB and CXC; methodology, SS, KED, YC, and CXC; formal analysis, SS, KED, and YC; investigation, SS, KED; writing—original draft preparation, SS; writing—review and editing, SS, KED, DB, and CXC; visualisation, SS; supervision, KED, DB, CXC; funding acquisition, DB and CXC. All authors have read and agreed to the published version of the manuscript.

## Acknowledgements

This project is supported by high-performance computing facilities at the National Computational Infrastructure (NCI) National Facility systems through the NCI Merit Allocation Scheme (Project d85) awarded to CXC, the University of Queensland Research Computing Centre, and computing facility at the Australian Centre for Ecogenomics, School of Chemistry and Molecular Biosciences at the University of Queensland.

