## Supplementary Figures S1 and S2 for "Intraspecies genomic divergence of coral algal symbionts shaped by gene duplication"

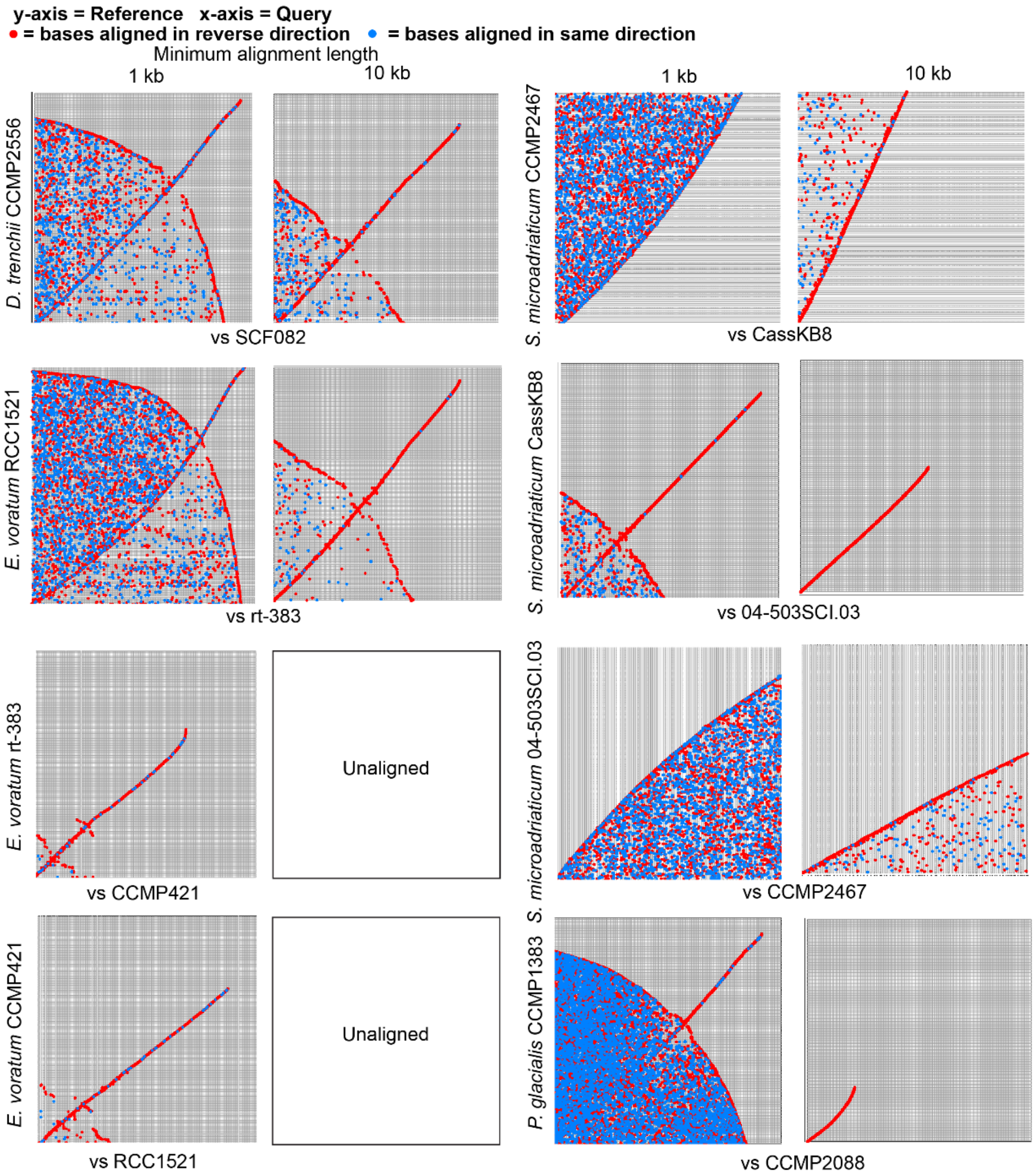


**Supplementary Figure S1**

Dot plots of intraspecies genome assemblies aligned to each other (minimum alignments 1 kb (grey), 10 kb (light grey)). The bases aligned between the reference (y-axis) versus the query (x-axis) genome assemblies, with alignments in the same direction represented in blue dots and reverse in red. No bases were aligned at 10 kb minimum lengths between *E. voratum* CCMP421 and RCC1521/rt-383.

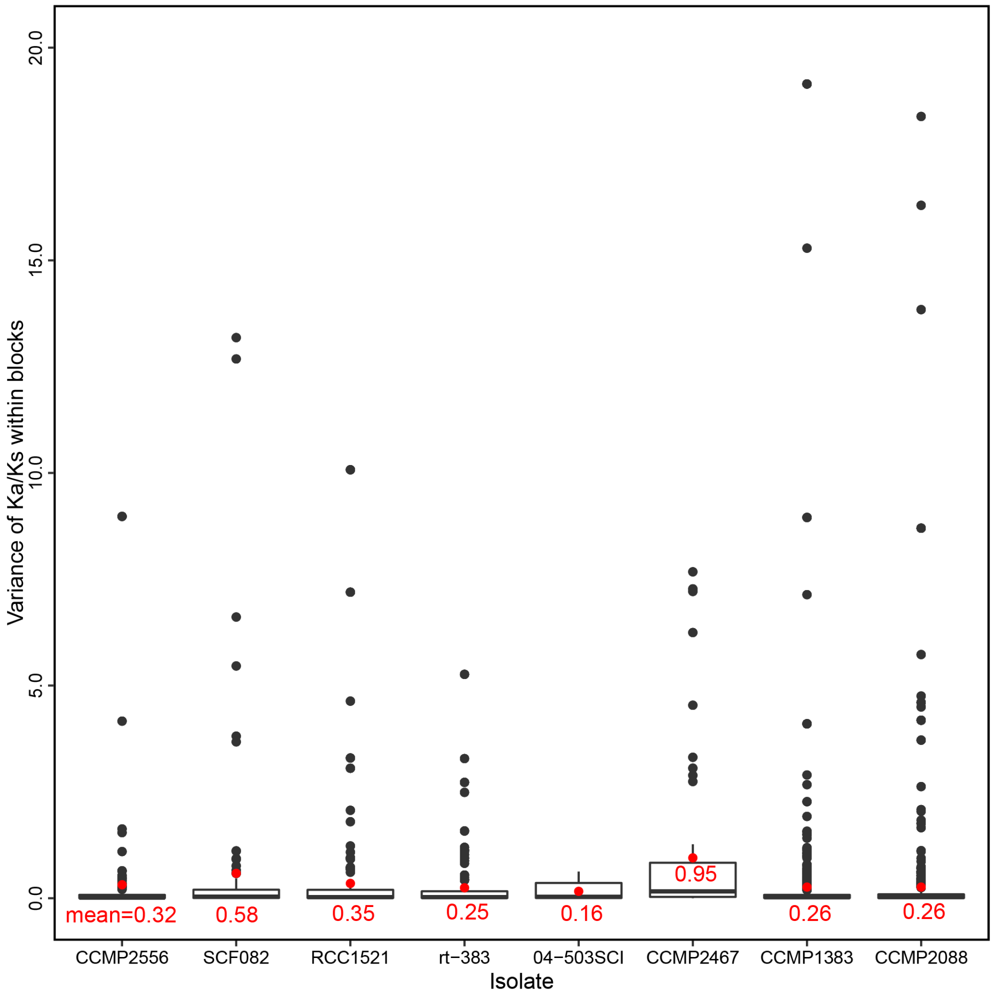


**Supplementary Figure S2**

The variance of mean $\omega$ within each TD block in each Suessiales isolate. The mean values are shown in red.
